## Supplemental Table 3 for "Adaptor protein supersaturation drives innate immune signaling and cell fate"

| **Plasmid name** | **Promoter** | **Resistance** | **Insert** | **Tag** | **ORF Sequence/Guide RNA** |
| --- | --- | --- | --- | --- | --- |
| lenti-CRISPR-FADD_g2 | EF-1α | Blasticidin | Cas9 |  | GAAGCTGGAGCGCGTGCAGAG |
| pCW57.1-PYCARD-mScarlet | tight TRE | Puromycin | PYCARD | (EA3R)4-mScarlet-I | MSGRARDAILDALENLTAEELKKFKLKLLSVPLREGYGRIPRGALLSMDALDLTDKLVSFYLETYGAELTANVLRDMGLQEMAGQLQAATHQGSGAAPAGIQAPPQSAAKPGLHFIDQHRAALIARVTNVEWLLDALYGKVLTDEQYQAVRAEPTNPSKMRKLFSFTPAWNWTCKDLLLQALRESQSYLVEDLERSEAAAREAAAREAAAREAAARINMVSKGEAVIKEFMRFKVHMEGSMNGHEFEIEGEGEGRPYEGTQTAKLKVTKGGPLPFSWDILSPQFMYGSRAFIKHPADIPDYYKQSFPEGFKWERVMNFEDGGAVTVTQDTSLEDGTLIYKVKLRGTNFPPDGPVMQKKTMGWEASTERLYPEDGVLKGDIKMALRLKDGGRYLADFKTTYKAKKPVQMPGAYNVDRKLDITSHNEDYTVVEQYERSEGRHSTGGMDELYK |
| pLV_AIM2-PYD-miRFP670-Cry2clust | EF-1α | Hygromycin | AIM2(PYD) | (EA3R)4-miRFP670nano-CRY2clust | MSESKYKEILLLTGLDNITDEELDRFKFFLSDEFNIATGKLHTANRIQVATLMIQNAGAVSAVMKTIRIFQKLNYMLLAKRLQEEKEKVDKQYGAEAAAREAAAREAAAREAAARIMANLDKMLNTTVTEVRQFLQVDRVCVFQFEEDYSGVVVVEAVDDRWISILKTQVRDRYFMETRGEEYSHGRYQAIADIYTANLTECYRDLLTQFQVRAILAVPILQGKKLWGLLVAHQLAAPRQWQTWEIDFLKQQAVVVGIAIQQSIRKPVATMKMDKKTIVWFRRDLRIEDNPALAAAAHEGSVFPVFIWCPEEEGQFYPGRASRWWMKQSLAHLSQSLKALGSDLTLIKTHNTISAILDCIRVTGATKVVFNHLYDPVSLVRDHTVKEKLVERGISVQSYNGDLLYEPWEIYCEKGKPFTSFNSYWKKCLDMSIESVMLPPPWRLMPITAAAEAIWACSIEELGLENEAEKPSNALLTRAWSPGWSNADKLLNEFIEKQLIDYAKNSKKVVGNSTSLLSPYLHFGEISVRHVFQCARMKQIIWARDKNSEGEESADLFLRGIGLREYSRYICFNFPFTHEQSLLSHLRFFPWDADVDKFKAWRQGRTGYPLVDAGMRELWATGWMHNRIRVIVSSFAVKFLLLPWKWGMKYFWDTLLDADLECDILGWQYISGSIPDGHELDRLDNPALQGAKYDPEGEYIRQWLPELARLPTEWIHHPWDAPLTVLKASGVELGTNYAKPIVDIDTARELLAKAISRTREAQIMIGAAARDPPDLDN |
| pLV_AIM2(F27G)-PYD-miRFP670-Cry2clust | EF-1α | Hygromycin | AIM2(F27G)(PYD) | (EA3R)4-miRFP670nano-CRY2clust | MSESKYKEILLLTGLDNITDEELDRFKGFLSDEFNIATGKLHTANRIQVATLMIQNAGAVSAVMKTIRIFQKLNYMLLAKRLQEEKEKVDKQYGAEAAAREAAAREAAAREAAARIMANLDKMLNTTVTEVRQFLQVDRVCVFQFEEDYSGVVVVEAVDDRWISILKTQVRDRYFMETRGEEYSHGRYQAIADIYTANLTECYRDLLTQFQVRAILAVPILQGKKLWGLLVAHQLAAPRQWQTWEIDFLKQQAVVVGIAIQQSIRKPVATMKMDKKTIVWFRRDLRIEDNPALAAAAHEGSVFPVFIWCPEEEGQFYPGRASRWWMKQSLAHLSQSLKALGSDLTLIKTHNTISAILDCIRVTGATKVVFNHLYDPVSLVRDHTVKEKLVERGISVQSYNGDLLYEPWEIYCEKGKPFTSFNSYWKKCLDMSIESVMLPPPWRLMPITAAAEAIWACSIEELGLENEAEKPSNALLTRAWSPGWSNADKLLNEFIEKQLIDYAKNSKKVVGNSTSLLSPYLHFGEISVRHVFQCARMKQIIWARDKNSEGEESADLFLRGIGLREYSRYICFNFPFTHEQSLLSHLRFFPWDADVDKFKAWRQGRTGYPLVDAGMRELWATGWMHNRIRVIVSSFAVKFLLLPWKWGMKYFWDTLLDADLECDILGWQYISGSIPDGHELDRLDNPALQGAKYDPEGEYIRQWLPELARLPTEWIHHPWDAPLTVLKASGVELGTNYAKPIVDIDTARELLAKAISRTREAQIMIGAAARDPPDLDN |
| pLV_APAF1(CARD)-miRFP670-Cry2clust | EF-1α | Hygromycin | APAF1(CARD) | (EA3R)4-miRFP670nano-CRY2clust | MDAKARNCLLQHREALEKDIKTSYIMDHMISDGFLTISEEEKVRNEPTQQQRAAMLIKMILKKDNDSYVSFYNALLHEGYKDLAALLHDGIPGAEAAAREAAAREAAAREAAARIMANLDKMLNTTVTEVRQFLQVDRVCVFQFEEDYSGVVVVEAVDDRWISILKTQVRDRYFMETRGEEYSHGRYQAIADIYTANLTECYRDLLTQFQVRAILAVPILQGKKLWGLLVAHQLAAPRQWQTWEIDFLKQQAVVVGIAIQQSIRKPVATMKMDKKTIVWFRRDLRIEDNPALAAAAHEGSVFPVFIWCPEEEGQFYPGRASRWWMKQSLAHLSQSLKALGSDLTLIKTHNTISAILDCIRVTGATKVVFNHLYDPVSLVRDHTVKEKLVERGISVQSYNGDLLYEPWEIYCEKGKPFTSFNSYWKKCLDMSIESVMLPPPWRLMPITAAAEAIWACSIEELGLENEAEKPSNALLTRAWSPGWSNADKLLNEFIEKQLIDYAKNSKKVVGNSTSLLSPYLHFGEISVRHVFQCARMKQIIWARDKNSEGEESADLFLRGIGLREYSRYICFNFPFTHEQSLLSHLRFFPWDADVDKFKAWRQGRTGYPLVDAGMRELWATGWMHNRIRVIVSSFAVKFLLLPWKWGMKYFWDTLLDADLECDILGWQYISGSIPDGHELDRLDNPALQGAKYDPEGEYIRQWLPELARLPTEWIHHPWDAPLTVLKASGVELGTNYAKPIVDIDTARELLAKAISRTREAQIMIGAAARDPPDLDN |
| pLV_NLRC4(CARD)-miRFP670-Cry2clust | EF-1α | Hygromycin | NLRC4(CARD) | (EA3R)4-miRFP670nano-CRY2clust | MNFIKDNSRALIQRMGMTVIKQITDDLFVWNVLNREEVNIICCEKVEQDAARGIIHMILKKGSESCNLFLKSLKEWNYPLFQDLNGQSLFGAEAAAREAAAREAAAREAAARIMANLDKMLNTTVTEVRQFLQVDRVCVFQFEEDYSGVVVVEAVDDRWISILKTQVRDRYFMETRGEEYSHGRYQAIADIYTANLTECYRDLLTQFQVRAILAVPILQGKKLWGLLVAHQLAAPRQWQTWEIDFLKQQAVVVGIAIQQSIRKPVATMKMDKKTIVWFRRDLRIEDNPALAAAAHEGSVFPVFIWCPEEEGQFYPGRASRWWMKQSLAHLSQSLKALGSDLTLIKTHNTISAILDCIRVTGATKVVFNHLYDPVSLVRDHTVKEKLVERGISVQSYNGDLLYEPWEIYCEKGKPFTSFNSYWKKCLDMSIESVMLPPPWRLMPITAAAEAIWACSIEELGLENEAEKPSNALLTRAWSPGWSNADKLLNEFIEKQLIDYAKNSKKVVGNSTSLLSPYLHFGEISVRHVFQCARMKQIIWARDKNSEGEESADLFLRGIGLREYSRYICFNFPFTHEQSLLSHLRFFPWDADVDKFKAWRQGRTGYPLVDAGMRELWATGWMHNRIRVIVSSFAVKFLLLPWKWGMKYFWDTLLDADLECDILGWQYISGSIPDGHELDRLDNPALQGAKYDPEGEYIRQWLPELARLPTEWIHHPWDAPLTVLKASGVELGTNYAKPIVDIDTARELLAKAISRTREAQIMIGAAARDPPDLDN |
| pCW57.1-CASP9-mScarlet | tight TRE | Puromycin | CASP9 | (EA3R)4-mScarlet-I | MDEADRRLLRRCRLRLVEELQVDQLWDALLSRELFRPHMIEDIQRAGSGSRRDQARQLIIDLETRGSQALPLFISCLEDTGQDMLASFLRTNRQAAKLSKPTLENLTPVVLRPEIRKPEVLRPETPRPVDIGSGGFGDVGALESLRGNADLAYILSMEPCGHCLIINNVNFCRESGLRTRTGSNIDCEKLRRRFSSLHFMVEVKGDLTAKKMVLALLELAQQDHGALDCCVVVILSHGCQASHLQFPGAVYGTDGCPVSVEKIVNIFNGTSCPSLGGKPKLFFIQACGGEQKDHGFEVASTSPEDESPGSNPEPDATPFQEGLRTFDQLDAISSLPTPSDIFVSYSTFPGFVSWRDPKSGSWYVETLDDIFEQWAHSEDLQSLLLRVANAVSVKGIYKQMPGCFNFLRKKLFFKTSGAEAAAREAAAREAAAREAAARINMVSKGEAVIKEFMRFKVHMEGSMNGHEFEIEGEGEGRPYEGTQTAKLKVTKGGPLPFSWDILSPQFMYGSRAFIKHPADIPDYYKQSFPEGFKWERVMNFEDGGAVTVTQDTSLEDGTLIYKVKLRGTNFPPDGPVMQKKTMGWEASTERLYPEDGVLKGDIKMALRLKDGGRYLADFKTTYKAKKPVQMPGAYNVDRKLDITSHNEDYTVVEQYERSEGRHSTGGMDELYK |
| pCW57.1-CASP1(CARD)-CASP9(93-416)-mScarlet | tight TRE | Puromycin | CASP1(CARD)-CASP9(93-416) | (EA3R)4-mScarlet-I | MADKVLKEKRKLFIRSMGEGTINGLLDELLQTRVLNKEEMEKVKRENATVMDKTRALIDSVIPKGAQACQICITYICEEDSYLAGTLGLSADRQAAKLSKPTLENLTPVVLRPEIRKPEVLRPETPRPVDIGSGGFGDVGALESLRGNADLAYILSMEPCGHCLIINNVNFCRESGLRTRTGSNIDCEKLRRRFSSLHFMVEVKGDLTAKKMVLALLELAQQDHGALDCCVVVILSHGCQASHLQFPGAVYGTDGCPVSVEKIVNIFNGTSCPSLGGKPKLFFIQACGGEQKDHGFEVASTSPEDESPGSNPEPDATPFQEGLRTFDQLDAISSLPTPSDIFVSYSTFPGFVSWRDPKSGSWYVETLDDIFEQWAHSEDLQSLLLRVANAVSVKGIYKQMPGCFNFLRKKLFFKTSGAEAAAREAAAREAAAREAAARINMVSKGEAVIKEFMRFKVHMEGSMNGHEFEIEGEGEGRPYEGTQTAKLKVTKGGPLPFSWDILSPQFMYGSRAFIKHPADIPDYYKQSFPEGFKWERVMNFEDGGAVTVTQDTSLEDGTLIYKVKLRGTNFPPDGPVMQKKTMGWEASTERLYPEDGVLKGDIKMALRLKDGGRYLADFKTTYKAKKPVQMPGAYNVDRKLDITSHNEDYTVVEQYERSEGRHSTGGMDELYK |
